## Supplementary materials for "Rapid and differential evolution of the venom composition of a parasitoid wasp depending on the host strain"

**Supplementary Figure S1: Biological model and experimental evolution protocol.** A. Interactions between *D. melanogaster* and *L. boulardi* strains. Host resistance to *L. boulardi* is only against the ISy line. Black arrow: encapsulated parasitoid egg inside a *D. melanogaster* larva. B. Design of the experimental evolution: ISm and ISy, ISm and ISy strains of *L. boulardi*; (R) and (S), resistant and susceptible host strains of *D. melanogaster*. F2, F6 and F10: the three analyzed generations of *L. boulardi*. C. Crossing tables showing the male and female expected genotypes in F1 and F2. "m" and "y" correspond to alleles from *L boulardi* ISm and ISy lines, respectively. The only assumption is mendelian inheritance.

**Supplementary Figure S2: Synthetic scheme of the analysis of the evolution of venom composition.** The global and specific approaches are described.

**Supplementary Figure S3: Complete mean intensity profile.** The graph shows (i) the median profile and part of its variability, (ii) the second derivative that was used to detect the peaks and (iii) the semi-automatically detected peaks. The solid and dotted black lines represent the weighted and unweighted median of intensities, respectively. Dotted grey lines are the weighted quartiles. The dotted blue line, the most informative, corresponds to the weighted median of the second derivatives used for the automatic detection of reference peaks. Colored vertical lines (blue, red and black) are the peak positions (local minima of the weighted median of second derivatives), and grey vertical lines are the borders of the peaks (local maxima of the weighted median of second derivatives). Black numbers below the horizontal 0 line are the ID of reference bands. Colored numbers (on grey dotted vertical lines) indicate the Rf coordinates of the borders of peaks. The color of these numbers corresponds to the color of the vertical line that indicates the position of the center of the peak to which borders coordinates refer. Colored numbers are positioned on four lines on the y axes, the two first ones corresponding to left borders, the others to right borders.

**Supplementary Figure S4: Details of the discriminant analysis**. The bars plot indicates the eigenvalues of the five discriminant axes. The three other plots show the position of individuals on the five discriminant axes identified. Individuals are grouped and colored according to the host strain and generation. (R), (S), resistant and susceptible host; subscript number, generation.

**Supplementary Table S1: Values of correlations of bands to discriminant axes.** Provided for the first two discriminant axes before and after partial correlation analysis. Cluster numbers are from the clustering analysis. Significance level: non-significant (n.s.); p-value > 0.05; * p-value < 0.05; ** p-value < 0.01; *** p-value <0.001. The two last columns indicate the significant correlations to axis 1 or 2 at the end of the analysis. Protein bands in bold represent the evolving protein bands at the end of the analysis. To be considered as an evolving protein band, the sign (+ or –) of the correlation had to be the same before and after the partial correlations, with a significance level lower than 0.05 before and after the partial correlations.

**Supplementary Methods: Procedure used to determine whether the LbGAP and LbSPN venom proteins were selected on the (R) and (S) host strains.** Calculation of the expected frequency of *lbspny* and *lbspnm* alleles and the [LbGAP] phenotype frequency. The file also contains the details of the script used for the simulations.

**Supplementary Figure S5: Statistical analysis of data from the specific approach for LbSPN and LbGAP proteins.** The genetic determinism is simple. Each panel shows the simulated null distributions of the modified chi$^2$ statistic describing the evolution of LbSPN (upper panels) and LbGAP (lower panels). For each protein, two hypotheses alternative to neutrality were considered: positive selection (left panel) or counter-selection (right panel). Vertical lines show the observed values of the modified chi$^2$ statistic computed using the populations reared either on the (R) (red line) or the (S) (blue line) host strain. Arrows highlight observations that deviate significantly from the null distribution.

**Supplementary Table S2: Modality comparisons for LbGAP2 quantity**. Multiple comparisons of means of each modality (host (S) or (R) and generation F2, F6 or F10) for the normalized LbGAP2 quantity of individuals. Tukey test implemented in the multcomp R package.

**Supplementary Figure S1**

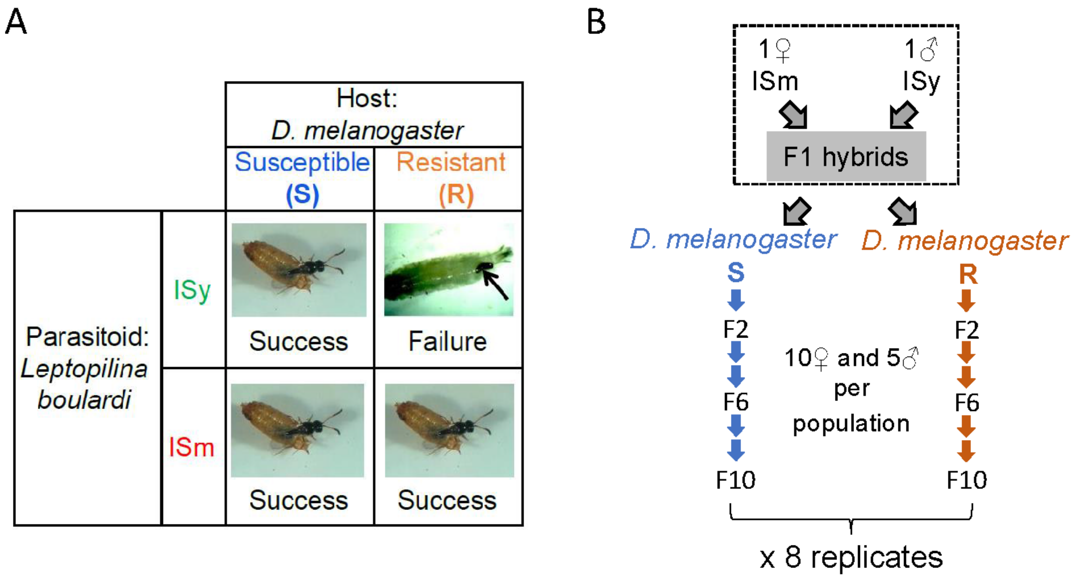


C

| ♀ ISm / ♂ ISy | ½ m | ½ m |
|---|---|---|
| y | **½ my** | **½ my** |

♀ F1: 100% my

| Virgin ♀ ISm | **½ m** | **½ m** |
|---|---|---|

♂ F1: 100% m

| ♀ F1 / ♂ F1 | ½ m | ½ y |
|---|---|---|
| m | **½ mm** | **½ my** |

♀ F2: 50% mm; 50% my

| Virgin ♀ F1 | **½ m** | **½ y** |
|---|---|---|

♂ F2: 50% m; 50% y

#### Supplementary Figure S2

### Analysis of the evolution of venom composition

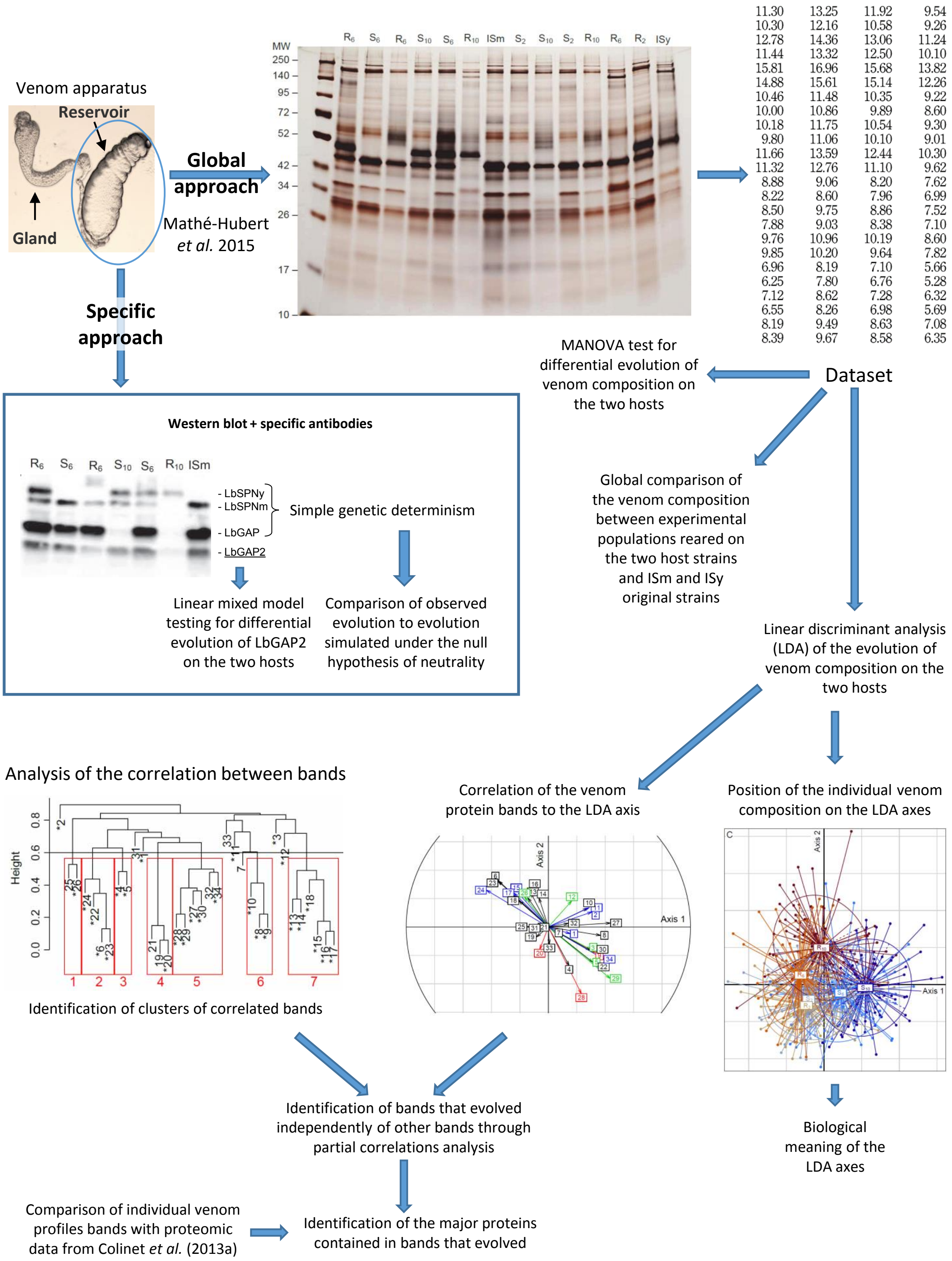

### Supplementary Figure S3

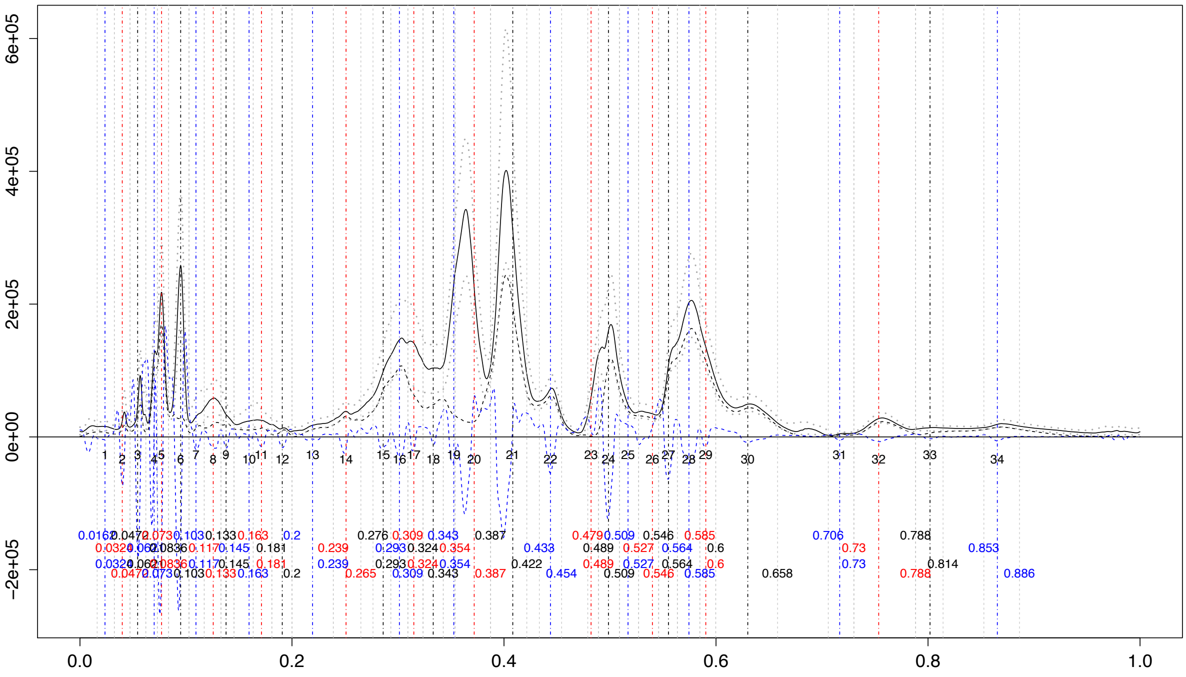

### Supplementary Figure S4

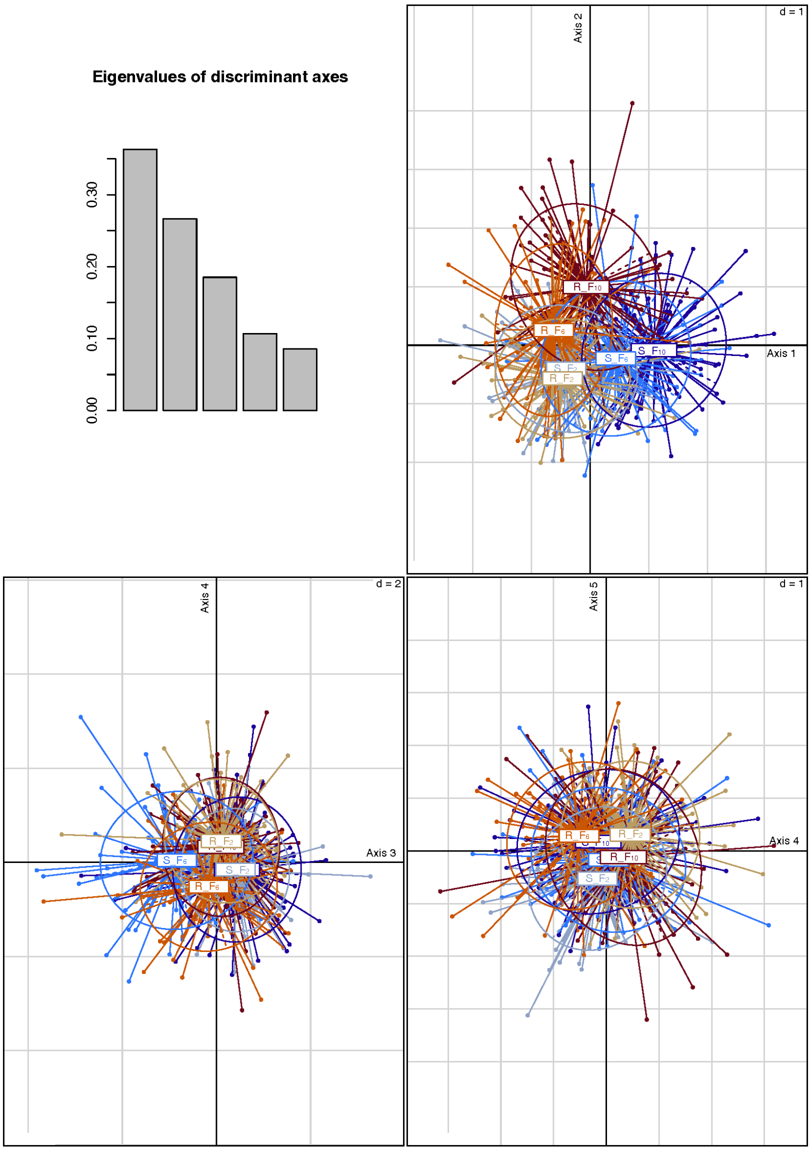

**Supplementary Table S1**

| Band | Cluster | Before partial correlation analysis | | | | After partial correlation analysis | | | | Summary of correlations | |
|---|---|---|---|---|---|---|---|---|---|---|---|
| | | Correlation with axis 1 | p-value associated | Correlation with axis 2 | p-value associated | Correlation with axis 1 | p-value associated | Correlation with axis 2 | p-value associated | Correlated with axis 1 | Correlated with axis 2 |
| **1** | - | **0.155** | **4.870e-02** | -0.047 | 1 | | | | | + | n.s. |
| **2** | - | **0.307** | **5.455e-10** | 0.106 | 1 | | | | | + | n.s. |
| **3** | - | **0.305** | **8.530e-10** | **-0.161** | **2.966e-02** | | | | | + | **-** |
| 7 | - | 0.048 | 1 | -0.042 | 1 | | | | | n.s. | n.s. |
| **11** | - | **0.288** | **1.149e-08** | 0.137 | 1.965e-01 | | | | | + | n.s. |
| **12** | - | **0.165** | **2.042e-02** | **0.185** | **3.548e-03** | | | | | + | + |
| 31 | - | -0.058 | 1 | -0.008 | 1 | | | | | n.s. | n.s. |
| 33 | - | 0.007 | 1 | -0.121 | 5.745e-01 | | | | | n.s. | n.s. |
| 25 | 1 | -0.151 | 6.657e-02 | 0.001 | 1 | | | | | n.s. | n.s. |
| **26** | 1 | **-0.167** | **1.722e-02** | **0.211** | **2.485e-04** | | | | | **-** | + |
| 6 | 2 | -0.351 | 2.161e-13 | 0.326 | 2.458e-11 | -0.056 | 1 | 0.053 | 1 | n.s. | n.s. |
| 22 | 2 | 0.352 | 2.067e-13 | -0.280 | 3.558e-08 | 0.147 | 5.220e-02 | -0.048 | 1 | n.s. | n.s. |
| 23 | 2 | -0.355 | 1.086e-13 | 0.325 | 2.958e-11 | 0.106 | 8.636e-01 | 0.048 | 1 | n.s. | n.s. |
| **24** | 2 | **-0.436** | **1.258e-21** | 0.260 | 6.416e-07 | **-0.273** | **7.174e-08** | 0.031 | 1 | **-** | n.s. |
| 4 | 3 | 0.146 | 9.754e-02 | -0.272 | 1.268e-07 | -0.012 | 1 | -0.129 | 0.193 | n.s. | n.s. |
| **5** | 3 | **0.313** | **2.016e-10** | **-0.248** | **2.900e-06** | **0.261** | **3.181e-07** | **-0.156** | **0.027** | + | **-** |
| 19 | 4 | -0.085 | 1 | -0.071 | 1 | | | | | n.s. | n.s. |
| **20** | 4 | -0.061 | 1 | **-0.158** | **3.852e-02** | | | | | n.s. | **-** |
| 21 | 4 | -0.005 | 1 | -0.002 | 1 | | | | | n.s. | n.s. |
| 27 | 5 | 0.438 | 8.712e-22 | 0.028 | 1 | 0.111 | 6.087e-01 | 0.369 | 2.770e-15 | n.s. | n.s. |
| **28** | 5 | 0.232 | 2.124e-05 | **-0.470** | **1.488e-25** | -0.172 | 6.627e-03 | **-0.213** | **1.217e-04** | n.s. | **-** |
| **29** | 5 | **0.435** | **1.673e-21** | **-0.363** | **2.454e-14** | **0.220** | **5.851e-05** | **-0.232** | **1.382e-05** | + | **-** |
| 30 | 5 | 0.297 | 2.901e-09 | -0.165 | 2.070e-02 | -0.080 | 1 | 0.053 | 1 | n.s. | n.s. |
| 32 | 5 | 0.141 | 1.377e-01 | 0.026 | 1 | | | | | n.s. | n.s. |
| **34** | 5 | **0.392** | **5.111e-17** | -0.227 | 3.804e-05 | **0.190** | **1.325e-03** | -0.146 | 5.694e-02 | + | n.s. |
| 8 | 6 | 0.369 | 7.120e-15 | -0.059 | 1 | 0.131 | 0.169 | 0.052 | 1 | n.s. | n.s. |
| **9** | 6 | 0.329 | 1.249e-11 | **-0.182** | **4.708e-03** | 0.041 | 1 | **-0.291** | **5.033e-09** | n.s. | **-** |
| 10 | 6 | 0.276 | 6.288e-08 | 0.133 | 2.519e-01 | 0.025 | 1 | 0.242 | 4.473e-06 | n.s. | n.s. |
| 13 | 7 | -0.145 | 1.037e-01 | 0.206 | 4.175e-04 | -0.020 | 1 | -8.806e-05 | 1 | n.s. | n.s. |
| 14 | 7 | -0.066 | 1 | 0.205 | 4.695e-04 | 0.077 | 1 | 7.119e-02 | 1 | n.s. | n.s. |
| **15** | 7 | **-0.212** | **2.264e-04** | 0.239 | 9.466e-06 | **-0.180** | **3.460e-03** | 6.763e-02 | 1 | **-** | n.s. |
| 16 | 7 | -0.123 | 4.885e-01 | 0.241 | 7.277e-06 | 0.262 | 3.399e-07 | 8.880e-03 | 1 | n.s. | n.s. |
| **17** | 7 | **-0.243** | **5.760e-06** | 0.240 | 8.844e-06 | **-0.199** | **5.127e-04** | -1.626e-02 | 1 | **-** | n.s. |
| 18 | 7 | -0.208 | 3.248e-04 | 0.1866 | 3.224e-03 | 0.010 | 1 | 4.795e-02 | 1 | n.s. | n.s. |

#### Supplementary Methods -

#### Procedure used to determine whether the LbGAP and LbSPN venom proteins were selected on the R and S host strains.

##### Summary of the procedure

The expected frequency of the ISm allele in our experimental populations was first computed considering a biallelic locus in an haplodiploid population, assuming (i) panmixia and (ii) neutrality (the tested $H_0$).

Then, a modified chi$^2$ statistics was used to summarize the deviation of experimental populations from the expectation.
This modification of the chi$^2$ statistics was devised to limit the effect of drift on the measured deviation.
Then, since the modified chi$^2$ statistics might not follow a chi$^2$ distribution anymore, the software simuPOP was used to simulate its null distribution.
The observed statistics describing the evolution on the R and S host strains was finally compared to this null distribution to get the p-value.

##### Computation of the expected frequency of the alleles / phenotype from ISm in the experimental populations

LbSPN is a co-dominant marker while LbGAP is a dominant marker. We thus used the allele frequencies for LbSPN and the phenotype frequency for LbGAP (sum of the frequencies of *lbgap*/*lbgap* and *lbgap*/*lbgap*$_y$ genotypes).

Notations:
M = ISm allele; Y = ISy allele
The null expectation for LbGAP is in yellow, and the null expectations for LbSPN are in green.

F0: fem = M/M; male = Y (Initial cross in the experimental evolution)
F1: fem = M/Y; male = M

| F1 → F2 females genotype | | Female gametes | |
|---|---|---|---|
| | | ½ M | ½ Y |
| Male gametes | M | MM | MY |

F2: fem = ½ MM ½ MY; male = ½ M ½ Y

| F2→F3 fem | ¾ M | ¼ Y |
|---|---|---|
| ½ M | MM | MY |
| ½ Y | MY | YY |

F3: fem = 3/8MM 1/2MY 1/8YY; male = 3/4 M ¼ Y

| F3→F4 fem | 5/8 M | 3/8 Y |
|---|---|---|
| ¾ M | MM | MY |
| ¼ Y | MY | YY |

F4: fem = 15/32MM 14/32 MY 3/32YY; male = 5/8 M 3/8 Y

| F4→F5 fem | 11/16 M | 5/16 Y |
|---|---|---|
| 5/8 M | MM | MY |
| 3/8 Y | MY | YY |

F5: fem = 55/128 MM 58/128 MY 15/128 YY; male = 11/16 M 5/16 Y

| F5→F6 fem | 21/32 M | 11/32 Y |
|---|---|---|
| 11/16 M | MM | MY |

| | | |
|---|---|---|
| 5/16 Y | MY | YY |

F6: fem = 231/512 MM 226/512 YM 55/512 YY; male = 21/32 M 11/32Y

LbGAP being dominant, frequencies for LbGAP = 231/512 + 226/512 = 457/512 and LbGAPy = 55/512

| F6→F7 fem | 43/64 M | 21/64 Y |
|---|---|---|
| 21/32 | MM | MY |
| 11/32 | MY | YY |

F7: fem = 903/2048 MM 914/2048 MY 231/2048 YY; male = 43/64 M 21/64 Y

| F7→F8 fem | 85/128 M | 43/128 Y |
|---|---|---|
| 43/64 M | MM | MY |
| 21/64 Y | MY | YY |

F8: fem = 3655/8192MM 3634/8192MY 903/8192 YY; male = 85/128 M 43/128 Y

| F8→F9 fem | 171/256 M | 85/256 Y |
|---|---|---|
| 85/128 M | MM | MY |
| 43/128 Y | MY | YY |

F9: fem = 14 535/32 768 MM 14 578/32 768 MY 3655/32 768 YY; male = 171/256 M 85/256 Y

| F9→F10 fem | 341/512 M | 171/512 Y |
|---|---|---|
| 171/256 M | MM | MY |
| 85/256 Y | MY | YY |

F10: fem = 58 311/131 072 MM 58 226/131 072 MY 14 535/131 072 YY; male = 341/512 M 171/512 Y

LbGAP being dominant, frequencies for LbGAP = 58311/131072+58226/131072=116537/131072 and LbGAPy = 14535/131

| F10→F11 fem | 683/1024 M | 341/1024 Y |
|---|---|---|
| 341/512 M | MM | MY |
| 171/512 Y | MY | YY |

This give the following expected values:

F6: LbSPNm: 43/64
LbGAP: 457/512

F10: LbSPNm: 683/1024
LbGAP: 116537/131072

**Formula of the modified chi² statistics using these four expectations**

The normal chi² statistics is:

$$\chi^2 = \sum \frac{(obs - exp)^2}{exp}$$

Here, *obs* and *exp* are respectively the observed and expected headcounts of the *lbspnm* allele frequencies or of the LbGAP phenotype in an experimental population at a given generation ($F_6$ and $F_{10}$), using either data for populations raised on the R host strain, or on the S host strain.

Drift will create deviation from the expected values in each replicate, in one direction or the other (some replicates being above the expectation and some below). Selection, on the opposite, will create deviation from the expected value in the same direction in all replicates. To measure the effects of selection rather than those of drift, we modified the chi² statistics as follows. This modification can be considered as a unilateral test of the chi² with the alternative hypothesis being H1 $obs < exp$ or $obs > exp$:

H1: $obs < exp$

$$\chi^2 = \sum \frac{(MIN\{obs - exp; 0\})^2}{exp}$$

H1: $obs > exp$

$$\chi^2 = \sum \frac{(MAX\{obs - exp; 0\})^2}{exp}$$

Because the modified chi² statistics might not follow anymore a chi² distribution, the software simuPOP was used to simulate the null distribution of these two statistics under the hypotheses of panmixia and neutrality for the simulated loci. This approach also tackles a problem we did not yet discussed. While we computed the statistics, we summed the divergences between observed and expected headcounts for each experimental population, even when they belong to the same replicate (for instance, one from the generation $F_6$ and one from the generation $F_{10}$) and are thus not independent. That would have been a problem if we had compared our observed chi² to the usual chi² distribution, but it is not here since the same formula was used to simulate the null distribution.

**Simulation of the null distribution of our statistics with the software simuPOP**

To obtain the null distribution of our statistics, the software simuPOP was used to simulate 20,000 times the neutral evolution of a bi-allelic locus evolving in the same conditions as in our experiment.
For each simulation, we simulated a biallelic locus evolving for 10 generations, in eight populations (our eight replicates), following an initial cross between males and females differing for their alleles to the considered genes (between the ISm and ISy strains). For each simulation, our summary statistic (the modified chi²) was computed in the same way as above, using the same expectation, and the number of *lbspnm* alleles for LbSPN, or of *lbgap* homozygous or heterozygous individual genotypes for LbGAP. As in the experiment, the populations' size was 10 females and 5 males for all generations.

P-values were obtained for each H1 hypothesis by comparing the observed statistics (modified chi²) to the corresponding null distribution. Since we tested the two H1 hypotheses ($obs < exp$ and $obs > exp$), the p-value was multiplied by two (Bonferroni correction).

**See below the script used to perform these simulations.**

```
# python3.5
###################################################################
###                     SET THE SIMULATION PARAMETERS                    ###
###################################################################
ISy = 1 # Value of the ISm and ISy alleles
ISm = 0
NsubPop = 8 # Number of replicats in the experiement

SimuleatedProt = 'LBSPN' # "LBSPN" or "LbGAP", LBSPN being co-dominant and LbGAP
dominant

Expected =[43/64     ,683/1024      ]# Expected frequencies of the LBSPN allele at F6
and F10
# Expected =[457/512   ,116537/131072 ]# Expected frequencies of the LbGAP allele at F6
and F10

H1 = 'inf' # 'sup' or 'inf': is the alternative hypothesis observed > (sup) expected, or
the opposite ?

AdresseOfSimulatedChi2 = "/supr/SimulatedChi2" # where to save the simulated modified
chi2?

NoSimulation = 20000

#############################################################
###                     PERFORM THE SIMULATION                     ###
#############################################################
import simuPOP as sim

if SimuleatedProt == 'LBSPN': # we are working with alleles (2n*N)
    Expected = [expect*20 for expect in Expected]

if SimuleatedProt == 'LbGAP': # we are working with individuals (N)
    Expected = [expect*10 for expect in Expected]

file = open(AdresseOfSimulatedChi2, "w")

def Chi2(pop): # function used to compute the modified chi2
    g1 = [None] * 8
    if SimuleatedProt == 'LBSPN':
        for r in range(NsubPop):
            g1[r] = sum([sum(x.genotype()) for x in pop.individuals(subPop=[r, 1])])
    if SimuleatedProt == 'LbGAP':
        for r in range(NsubPop):
            g1[r] = sum([max(x.genotype()) for x in pop.individuals(subPop=[r,1])])
    if pop.dvars().Generation == 6:
        if H1 == 'sup':
            chi2 = sum([ (max([(g-Expected[0]),0])**2)/Expected[0] for g in g1])
        if H1 == 'inf':
            chi2 = sum([ (min([(g-Expected[0]),0])**2)/Expected[0] for g in g1])
        pop.dvars().chi2 = chi2
    if pop.dvars().Generation == 10:
        if H1 == 'sup':
            chi2 = sum([ (max([(g-Expected[1]),0])**2)/Expected[1] for g in g1])
        if H1 == 'inf':
            chi2 = sum([ (min([(g-Expected[1]),0])**2)/Expected[1] for g in g1])
        pop.dvars().chi2 =+ chi2
        file.write(str(pop.dvars().chi2) + '\n')
    return True
```

```
for rep in range(NoSimulation):
    pop = sim.Population(size=[15]*NsubPop, loci=1, ploidy=sim.HAPLODIPLOID
    pop.setVirtualSplitter(sim.CombinedSplitter(splitters=[
        sim.SexSplitter(),

sim.ProductSplitter(splitters=[sim.SexSplitter(),sim.GenotypeSplitter(loci=[0],alleles=[
[0],[1]]) ] )
    ]))
    #pop.subPopName([0, 0])
    #   'Male'
    #pop.subPopName([0, 1])
    #   'Female'
    pop.dvars().chi2 = None
    pop.dvars().Generation = 0 # We start by performing the initial cross between ISm
and ISy
    pop.evolve(  # describeEvolProcess
        initOps=[
            sim.InitSex(sex=[sim.MALE]*5 +[sim.FEMALE]*10),
            sim.InitGenotype(genotype=[ISm, ISm],subPops=[(sim.ALL_AVAIL,1)]),
            sim.InitGenotype(genotype=[ISy, ISy],subPops=[(sim.ALL_AVAIL,0)])
        ],
        preOps=[sim.PyOperator(Chi2)],
        matingScheme=sim.HaplodiploidMating(
sexMode=(sim.GLOBAL_SEQUENCE_OF_SEX,sim.MALE,sim.FEMALE,sim.FEMALE)
                                                 ,ops=sim.HaplodiploidGenoTransmitter()),
    postOps = [sim.PyExec('Generation += 1')
               ]
        ,gen=11)# we start with F0


file.close()
```

### Supplementary Figure S5

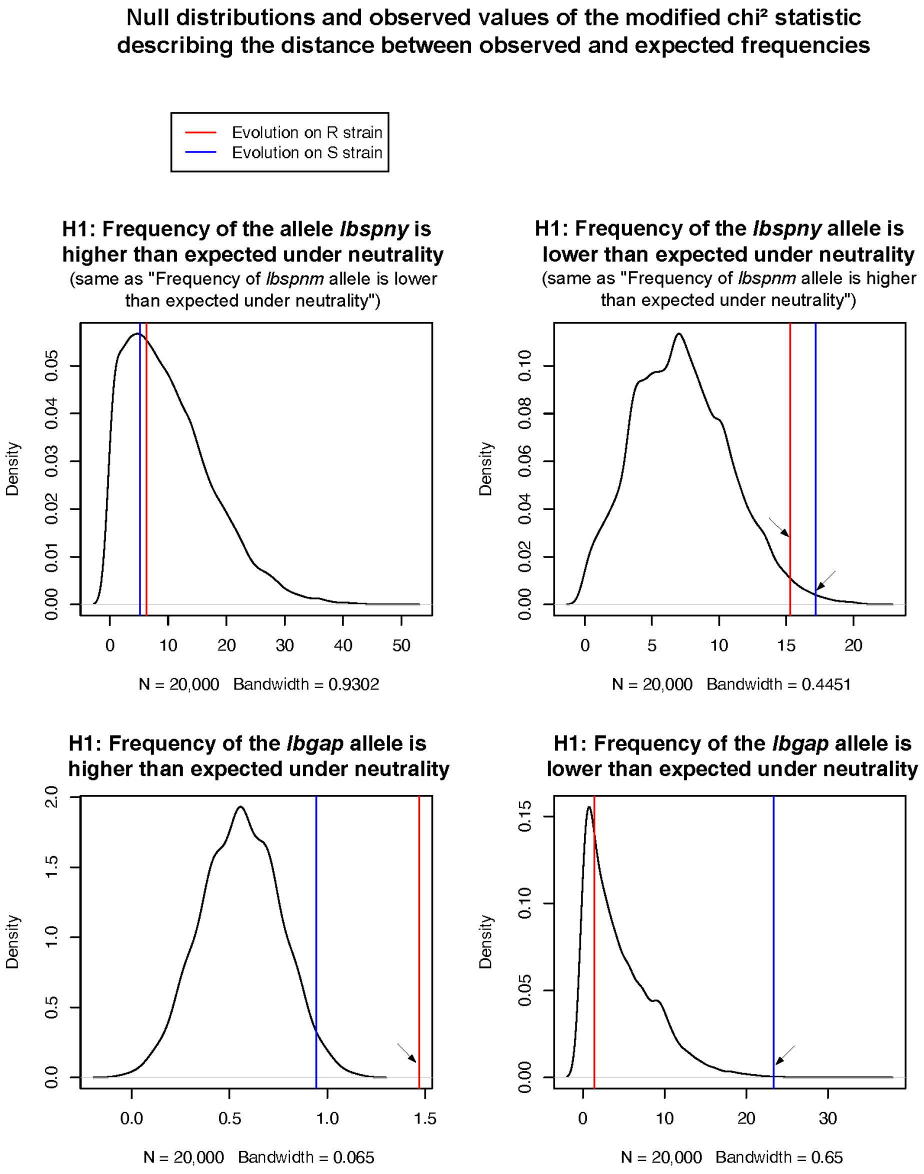

**Supplementary Table S2.**

| $H_0$ | Estimate | Std. Error | z value | Pr(>\|z\|) |
|---|---|---|---|---|
| $R\_F_2 - S\_F_2 = 0$ | 0.077464 | 0.102156 | 0.758 | 0.9739 |
| $S\_F_6 - S\_F_2 = 0$ | 0.004686 | 0.091679 | 0.051 | 1.0000 |
| $R\_F_6 - S\_F_2 = 0$ | -0.285879 | 0.102144 | -2.799 | 0.0568. |
| $S\_F_{10} - S\_F_2 = 0$ | -0.217993 | 0.092019 | -2.369 | 0.1653 |
| $R\_F_{10} - S\_F_2 = 0$ | -0.505383 | 0.102403 | -4.935 | **<0.001** *** |
| $S\_F_6 - R\_F_2 = 0$ | -0.072778 | 0.100324 | -0.725 | 0.9785 |
| $R\_F_6 - R\_F_2 = 0$ | -0.363343 | 0.090282 | -4.025 | **<0.001** *** |
| $S\_F_{10} - R\_F_2 = 0$ | -0.295457 | 0.100579 | -2.938 | 0.0385 * |
| $R\_F_{10} - R\_F_2 = 0$ | -0.582846 | 0.090544 | -6.437 | <0.001 *** |
| $R\_F_6 - S\_F_6 = 0$ | -0.290565 | 0.100325 | -2.896 | 0.0431 * |
| $S\_F_{10} - S\_F_6 = 0$ | -0.222679 | 0.089972 | -2.475 | 0.1300 |
| $R\_F_{10} - S\_F_6 = 0$ | -0.510069 | 0.100583 | -5.071 | **<0.001** *** |
| $S\_F_{10} - R\_F_6 = 0$ | 0.067886 | 0.100579 | 0.675 | 0.9844 |
| $R\_F_{10} - R\_F_6 = 0$ | -0.219504 | 0.090555 | -2.424 | 0.1461 |
| $R\_F_{10} - S\_F_{10} = 0$ | 0.287390 | 0.100839 | -2.850 | 0.0494 * |
